# RNA secondary structure regulates fragments’ adsorption onto flat substrates

**DOI:** 10.1101/2021.08.31.458432

**Authors:** Simón Poblete, Anže Božič, Matej Kanduč, Rudolf Podgornik, Horacio V. Guzman

## Abstract

RNA is a functionally rich molecule with multilevel, hierarchical structures whose role in the adsorption to molecular substrates is only beginning to be elucidated. Here, we introduce a multiscale simulation approach that combines a tractable coarse-grained RNA structural model with an interaction potential of a structureless flat adsorbing substrate. Within this approach, we study the specific role of stem-hairpin and multibranch RNA secondary structure motifs on its adsorption phenomenology. Our findings identify a dual regime of adsorption for short RNA fragments with and without secondary structure, and underline the adsorption efficiency in both cases as a function of the surface interaction strength. The observed behavior results from an interplay between the number of contacts formed at the surface and the conformational entropy of the RNA molecule. The adsorption phenomenology of RNA seems to persist also for much longer RNAs as qualitatively observed by comparing the trends of our simulations with a theoretical approach based on an ideal semiflexible polymer chain.

## Introduction

In the past decades, the ability of ribonucleic acid (RNA) to influence biological processes occurring inside living cells^1^ has generated a lot of interest and has provided a driving force for innovative strategies in ambitious nanomedical applications.^2–6^ RNA is a flexible polyelectrolyte with highly adaptable conformations, and with self-associating base pairs creates a variety of complex structural motifs, which sets it apart from the commonly more rigid and structurally much less diverse DNA.^7–9^ Unlike proteins, RNA acquires its structure in a hierarchical way, first assuming a secondary structure—a pattern of base pairs—followed by the formation of a three dimensional tertiary structure.^10–15^ While RNA differs fundamentally from DNA with its pervasive, stable double-stranded form, its self-association bears some similarity with protein folding, though the structural motifs present in RNA tend to be in general much softer and less globular.^16^ This conformational softness furthermore implies that RNA might structurally respond to the presence of interactions with other macromolecular substrates sharing its biological environment. An important mode of these interactions is related to RNA adsorption to proteinaceous substrates, such as the capsid shells of viruses, where the RNA-capsid interactions are important for the efficiency of virion assembly and nanoparticle stability.^17–19^ In this context, the fundamental question is whether the self-assembled RNA conformation is modified as a result of the adsorption or, equivalently, whether different RNA structural motifs modify its adsorption phenomenology.

From a fundamental polymer theory point of view,^20–23^ molecular simulations have offered important insights into the adsorption of semiflexible macromolecules to a molecular substrate,^24^ while the adsorption of macromolecules with either annealed or quenched internal structure onto a molecular substrate remains much less understood. The latter problem is particularly relevant in the context of RNA-virus and RNA-nanoparticle assembly phenomena,^25–32^ where the soft, malleable RNA structure can respond to the adsorption process. The shortage of theoretical conceptualization and prediction makes it difficult to convert the observed RNA adsorption phenomenology into a robust parametrization of the underlying adsorption interaction potential and consequently modify and/or control the RNA-substrate interactions. Such insight into the adsorption characteristics would be especially valuable for the optimization and control of RNA assembly into carrier vesicles or virus-like nanoparticles for efficient RNA-cargo delivery,^33–36^ which could potentially speed up high-throughput RNA nanocarrier fabrication for applications in nanomedicine.

The lack of a solid understanding of the connection between self-assembled structures of biopolymers such as RNA,^37,38^ induced by specific internucleotide interactions, and their modification as a result of the adsorption process, induced by less specific interactions with the adsorbing substrate, is a challenge whose general aspects we aim to address in this work. We have performed an extensive study of RNA-substrate interaction using a tractable multiscale^39–41^ model to understand the adsorption mechanisms of RNA in proximity to a flat, featureless model of an adsorbing substrate. In order to elucidate the role of the secondary structure on the adsorption mechanisms of RNA, we performed coarse-grained molecular simulations of RNA fragments extracted from the Satellite Tobacco Mosaic Virus (STMV) genome.^42^ One particular question we address is the role of different soft RNA secondary structure motifs in the adsorption phenomenology and the connection between the inherent RNA structure and the structure imposed by the adsorption itself. To this end, we compare the adsorption phenomenology of RNA fragments in two distinct configurations, with and without secondary structure. The model interaction potential between the adsorbing substrate and the proximal structured or unstructured RNA molecule is allowed to vary in a noninvasive manner so that the original secondary structures, if they exist, remain fixed.

We identify a transition between two interaction regimes for structured and unstructured RNA as the attractive substrate strength is varied. For structured RNA (with secondary structure), base-paired regions are preferentially adsorbed at lower surface interaction strengths when compared to the unstructured RNA. For unstructured RNA (without secondary structure), the opposite is true, and they are preferentially adsorbed at larger adsorption strengths compared to structured RNA. While it is difficult to scale up our simulations to significantly longer RNA sequences, we do simulate the adsorption behavior for different RNA sizes and compare their adsorption behavior to the ground state behavior of ideal polymer chains with the same adsorption potential. Finally, we discuss the possible role that the different adsorption regimes could play in the nanomedicine applications that can be complemented with our simulations.

## System and Model Description

### RNA fragments

Our study focuses on three short RNA fragments with different basic secondary structure motifs (shown in Figure 1): a short hairpin (S1), a large hairpin including several bulges (S2), and a two-hairpin multibranch structure of a similar length (S3). All three RNA fragments were extracted from previous experimental studies on the genome of STMV, whose secondary structures has been determined by SHAPE chemical probing method.^43,44^ It is important to remark that the three fragments of this work represent archetypal structures of the entire STMV genome, as previously obtained by chemical probing methods.^42^ Our coarse-grained model generates a 3D representation of the three RNA fragments and supports secondary structure restraints based on primary sequence and interactions deconvolved from an X-ray structure database (see Methods for details).

**Figure 1:**
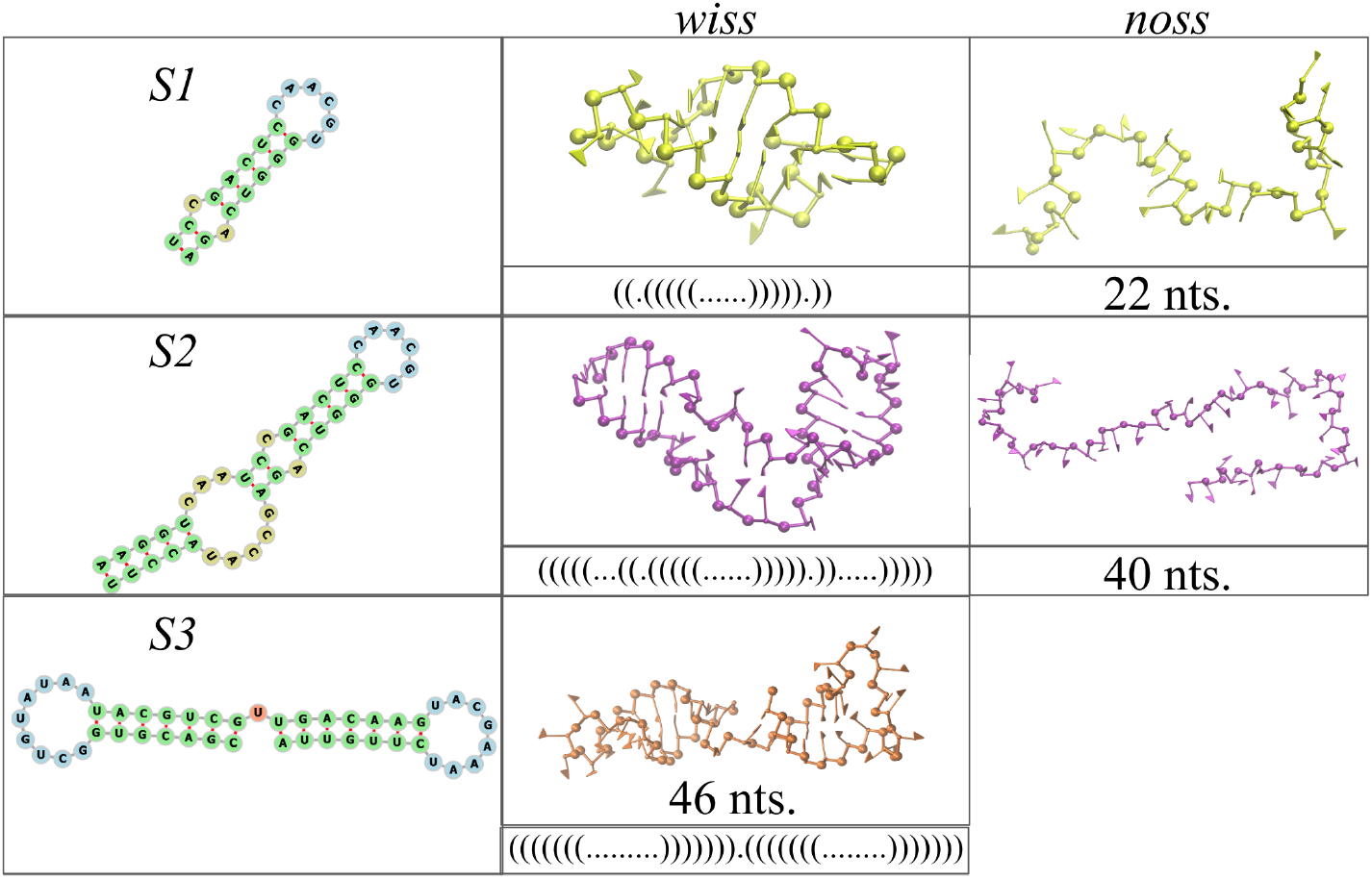
Secondary structure (left column), 3D model with secondary structure (middle column), and 3D model without secondary structure (right column) for the three RNA fragments used in this study (S1, S2, and S3).

In order to explore the role of the secondary structure in RNA adsorption to a model substrate, we consider representative structured (“wiss”—with secondary structure) and unstructured (“noss”—no secondary structure) versions of the first two fragments, S1 and S2, while the third fragment complements the set by introducing a multibranch secondary structure (as illustrated in Figure 1). We did not consider the unstructured counterpart of the third fragment, as it is of a similar length to the second one (S2).

### Model RNA-substrate interaction

We aim to investigate the general role of RNA secondary structure in adsorption processes. To that purpose, we model the substrate as a flat, featureless surface. Excluding topographical and molecular features of the substrate, which could possibly intervene in the interpretation of the results, allows us to isolate the influence of the RNA secondary structure in its adsorption. We model the attraction of the RNA phosphate groups to the adsorbing substrate by a Debye-Hückel-like interaction potential, which can be rationalized as stemming from the electrostatic interactions between the dissociated RNA phosphates and the substrate charges.^45^ Combined with a generic short-range repulsive term, the surface potential acting on the RNA then assumes the form

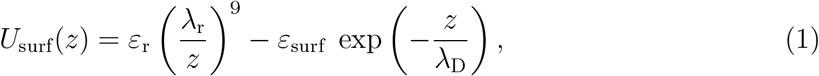

where *ε*_r_ = 1 *k*_B_*T*, λ_r_ = 0.1 nm is the distance of activation of the repulsive Lennard-Jones term (well below the size of the RNA phosphates ≈ 0.3 nm, as defined in our model) and λ_D_ = 1 nm, considering RNA under typical physiological conditions as described in Refs. 46,47. The strength of the attractive potential, *ε*_surf_, is varied in the range between 0.44 *k*_B_*T* and 1.78 *k*_B_*T*. For comparison, we also considered the scenario in which the RNA interacts with the substrate via the Mie 9-3 potential.^48^ The latter Lennard-Jones-based potential can be rationalized as stemming only from the van der Waals interactions. The results are reported in the Supplementary Information.

Temperature and structural restraints of the system are chosen in such that they do not destroy the secondary structure imposed, corresponding to a scenario below the melting temperature (for details, see the Supplementary Information). Figure 2 shows a schematic representation of the most prominent features of a simple RNA hairpin with (right) and without (left) the secondary structure, together with the action of the surface interaction potential and the center-of-mass coordinate *d*_CM_.

**Figure 2:**
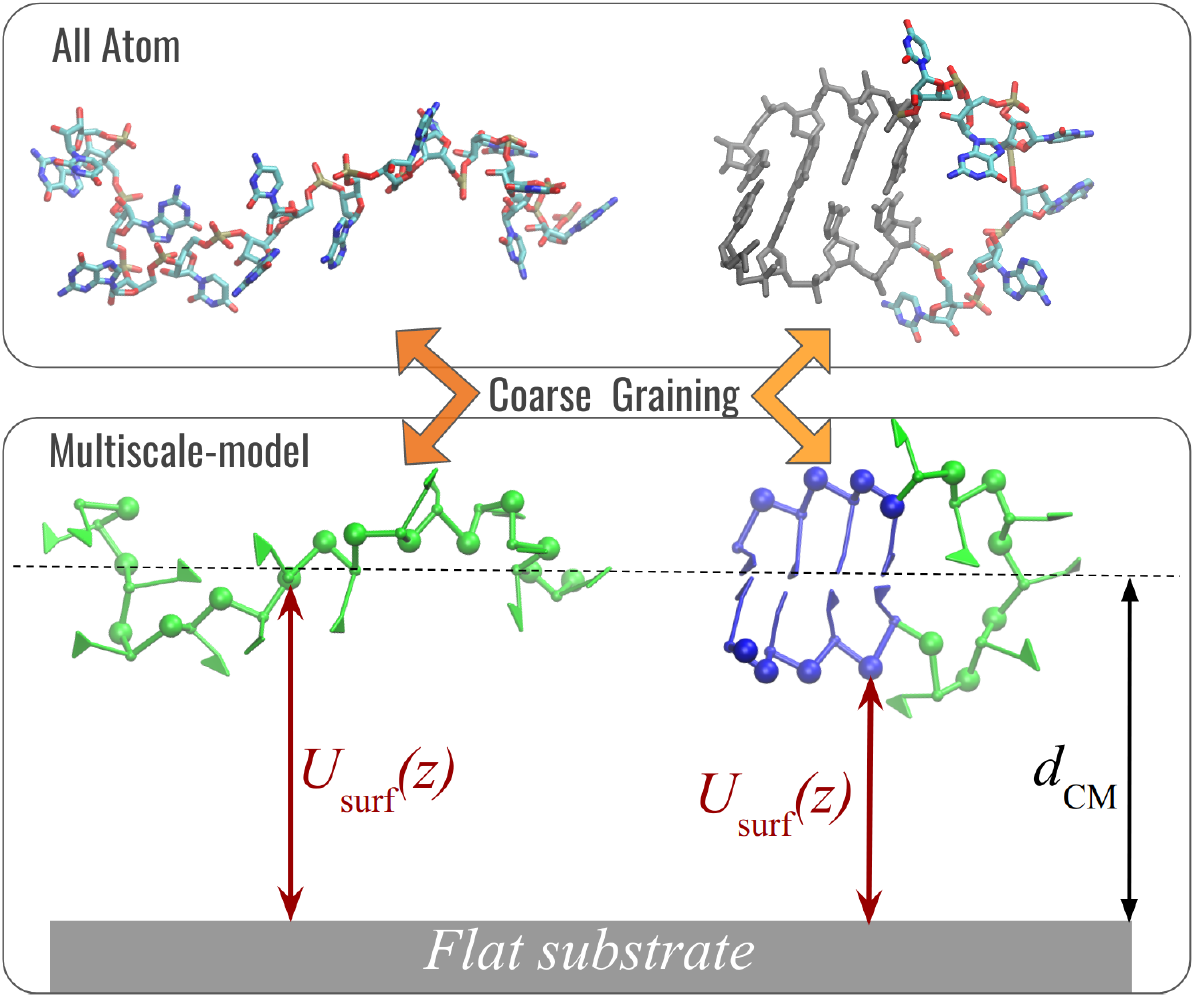
Two representative snapshots of a structured hairpin RNA fragment (right) and an unstructured (left) in All Atom (top) and multiscale representation (bottom). Coarsegrained nucleotides are comprised of five beads: a triangle in the bases, made of three sites, a bead representing the sugar, and a bead representing the phosphate. RNA phosphates are subjected to the RNA-substrate interaction potential *U*_surf_, Equation 1. The schematic hairpin shows the base-paired nucleotides in blue and the unpaired ones in green. All-atom representation is sketched here as part of the illustration of the coarse-graining within the multiscale model.

## Results and Discussion

### Small hairpin (RNA fragment S1)

The first RNA motif we address is a hairpin with a small bulge, comprising 22 nucleotides with a total of 7 base pairs, as shown in Figure 1. The behavior of the calculated potential of mean force (PMF) as a function of the distance of the RNA center-of-mass from the substrate is plotted in Figure 3. The PMF was calculated by averaging different conformations of the flexible regions, as well as orientations of the base-paired fragments; see also Supplementary Information. Figure 3 shows the results for both the structured (A) and the unstructured (B) S1 fragments. The PMF has the form of an attractive well, with a longer range for the unstructured fragment, and with a minimum whose location and depth depend on the strength of the RNA-substrate attraction (controlled by *ε*_surf_; see Figure 3(B)). Given the restraints of its secondary structure, the S1-wiss fragment cannot get closer than 0.9 nm from the substrate, while the single-stranded fragment can deform and adsorb more efficiently if the attraction to the substrate is strong enough.

**Figure 3:**
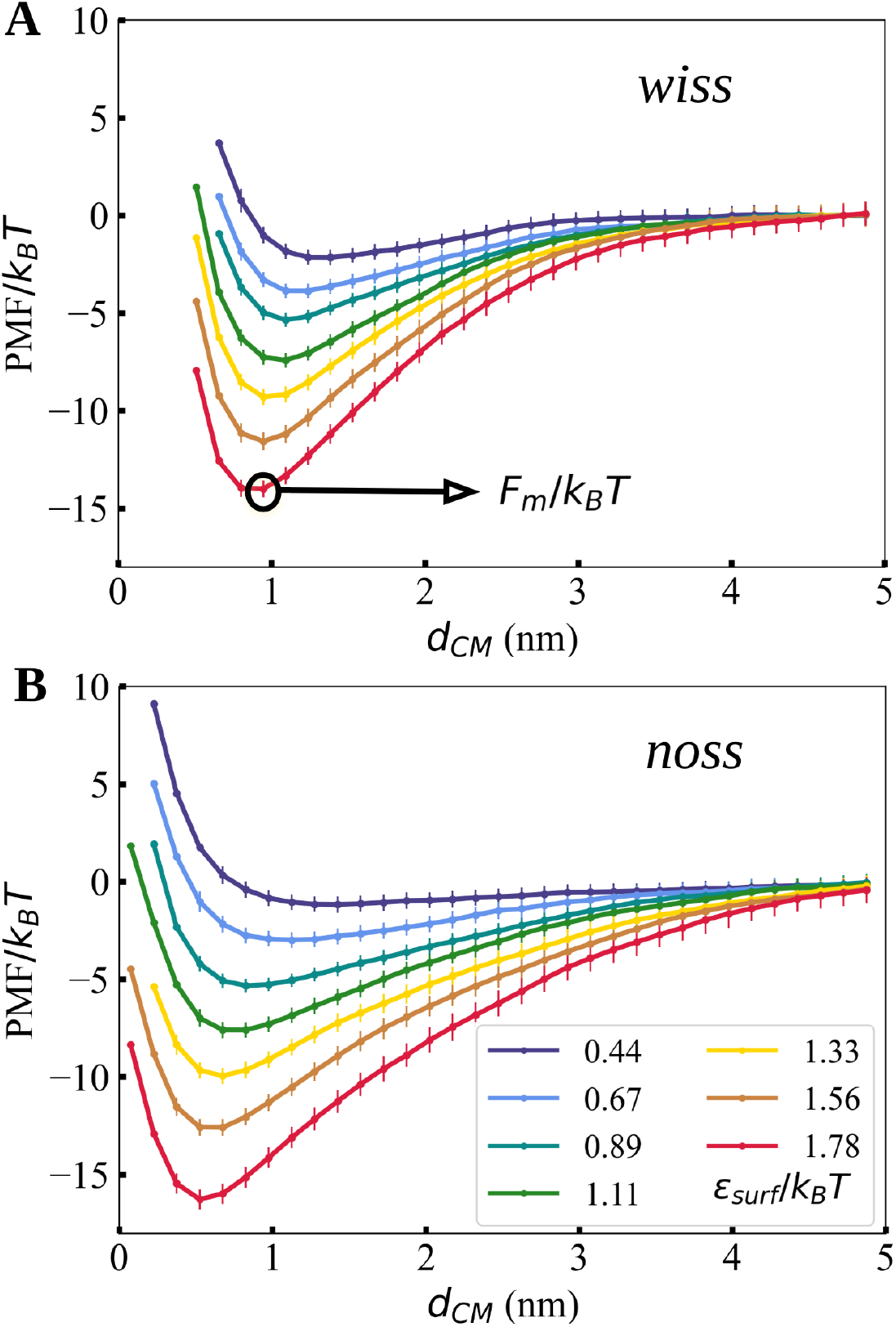
Potential of mean force as a function of the distance between the substrate and the center-of-mass of RNA fragment S1 **(A)** with and **(B)** without the secondary structure, shown for representative values of the attraction strength *ε*_surf_ in normalized units.

Figure 4(A) shows adsorption free energy *F*_m_ (defined as the minimum of PMF) as a function of the surface attraction strength *ε*_surf_, for both structured and unstructured RNA fragments. Two regimes are immediately identifiable: the first for *ε*_surf_ < 0.89*k*_B_*T*, where the fragment with secondary structure adsorbs more strongly than the unstructured one (regime I), while for *ε*_surf_ > 0.89*k*_B_*T*, the unstructured fragment is the one exhibiting a stronger adsorption free energy (regime II).

**Figure 4:**
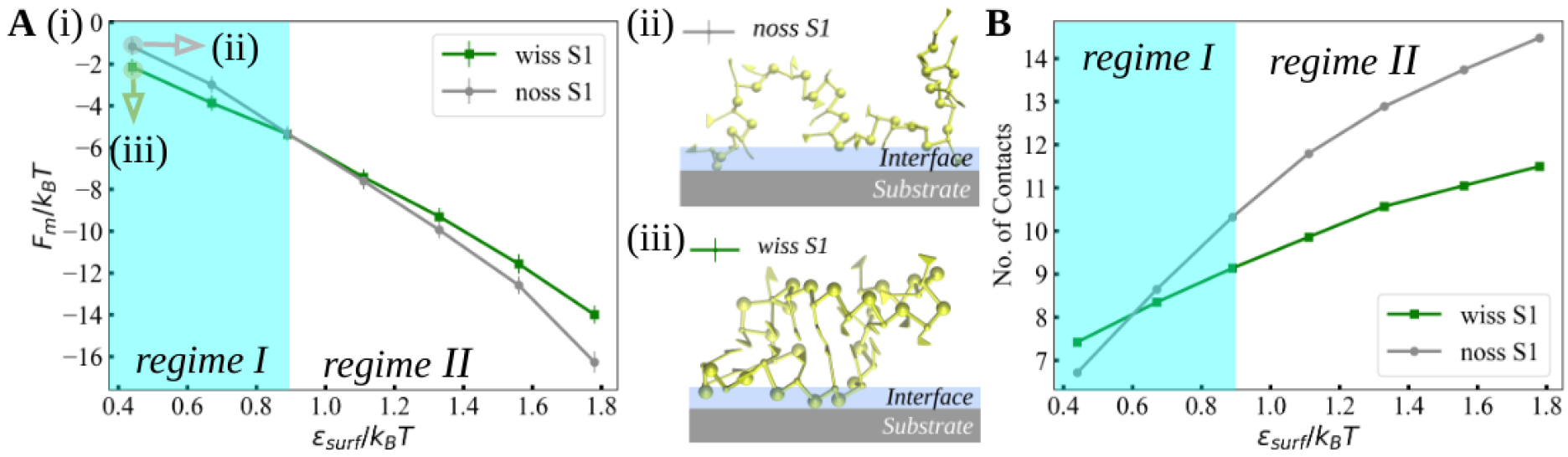
**(A)** Adsorption free energy as a function of the substrate attraction strength, *ε*_surf_/*k*_B_*T*, for RNA fragments S1-wiss and S1-noss. For a weak attraction of *ε*_surf_/*k*_B_*T* = 0.44, we show simulation snapshots of the (ii) unstructured and (iii) structured fragments. The illustrated interface layer is a schematic definition of a distance slightly thicker than the diameter of phosphates. **(B)** Number of contacts as a function of *ε*_surf_/*k*_B_*T* for RNA fragments S1-wiss and S1-noss, calculated via Equation 2.

An interesting question is how general is the existence of the two adsorption regimes. To provide a rough answer, we performed simulations using a different type of attractive potential, namely, a Lennard-Jones-based potential for a planar wall (Mie 9-3 potential). The outcomes of this much shorter-ranged potential are qualitatively the same as for the electrostatically-driven, longer-ranged Debye-Hückel potential (see Figures S1 and S2 in Supplementary Information). This suggests that the existence of the two adsorption regimes is not specific to the Debye-Hückel potential but may be considered a more general phenomenon, common to other substrate architectures as well.

We quantify the number of contacts of RNA with the surface based on the normalized energy of the RNA-substrate interaction:

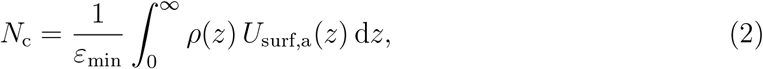

where *U*_surf,a_(*z*) = min{0, *U*_surf_(*z*)}, *ε*_min_ is the minimum of the surface potential, and *ρ*(*z*) is the monomer distribution for bound conformations normalized as 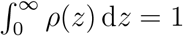 (for details and description, see Methods and Figure S3 in Supplementary Information). Equation 2 gives the exact contact fraction for a square-well surface potential, which is well-studied in the literature.^23^

As shown in Figure 4(B), on weakly attractive surfaces, the unstructured fragment forms fewer contacts than the structured one. This behavior changes at *ε*_surf_ = 0.6 *k*_B_*T*, which is below the point where the adsorption free energies *F*_m_ coincide (Figure 4(A)). Nevertheless, this behavior suggests that the adsorption free energy is influenced by the number of contacts formed with the surface and also by the conformation entropy of the RNA molecules. At high interaction strengths, *i.e*., *ε*_surf_ > 0.6 *k*_B_*T*, the unstructured fragment creates more contacts with the substrate than the structured one (Figure 4(B)). This result implies that the secondary structure of the fragment controls the number of contacts that can be made with the substrate. Furthermore, it suggests that the unstructured fragment can exhibit a saturated adsorption with all monomers being in contact with the substrate, like an RNA “landing pad.” Note also that in our simulations the secondary structure, if it exists, remains fixed. Monomer distributions for RNA fragments S2 and and S3 are shown in Figure S4 in Supplementary Information, and the number of contacts for the two fragments are shown in Figure S5 in Supplementary Information. For fragment S2, the intersection between the wiss and noss occurs at slightly higher attraction strength than for S1, namely at *ε*_surf_/*k*_B_*T* = 0.7. The exact value of the intersection point depends on the fraction of structured domains. Moreover, hairpins, internal loops or junctions might contribute in a different manner to the phosphate distribution from the surface and the contact fraction.

We obtain further insight into the structural configurations of the RNA molecules by looking at the parallel and perpendicular contributions to the radius of gyration (defined in Supplementary Information). In Figure 5(A), the ratio between the normal 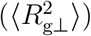 and parallel 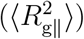 contributions to the radius of gyration of fragment S1 in contact with the surface is shown to decrease monotonically as the substrate attraction gets stronger. A slower decrease in 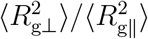 is observed in the structurally constrained fragment given its flexible loop, while for unstructured RNA the values fall well below 0.32, in agreement with theoretical predictions for semiflexible polymer chains.^23^ Representative snapshots of fragment S1 are shown as a side view in Figures 5(B) and (C) for the structured and unstructured cases, respectively. Whereby, we also observe more flattened conformations for the unstructured fragment and for stronger substrate attractions, which is a consequence of an increased formation of contacts in the unstructured RNA fragments and their consequent entropy loss. Figures 5(D) and (E) show the top view of the same fragments, with a discernible extension in the *xy*-plane for the unstructured fragment and a reorganization of structural constrains for the structured fragment. The flattening of the unstructured RNA fragments reflects also in the decrease of the ratio between the normal and parallel radii of gyration with increasing surface attraction strength (shown in Figure 5(A)), and the fact that the parallel contribution is less sensitive to *ε*_surf_ than its normal counterpart (see Tables S1 and S2 in the Supplementary Information).

**Figure 5:**
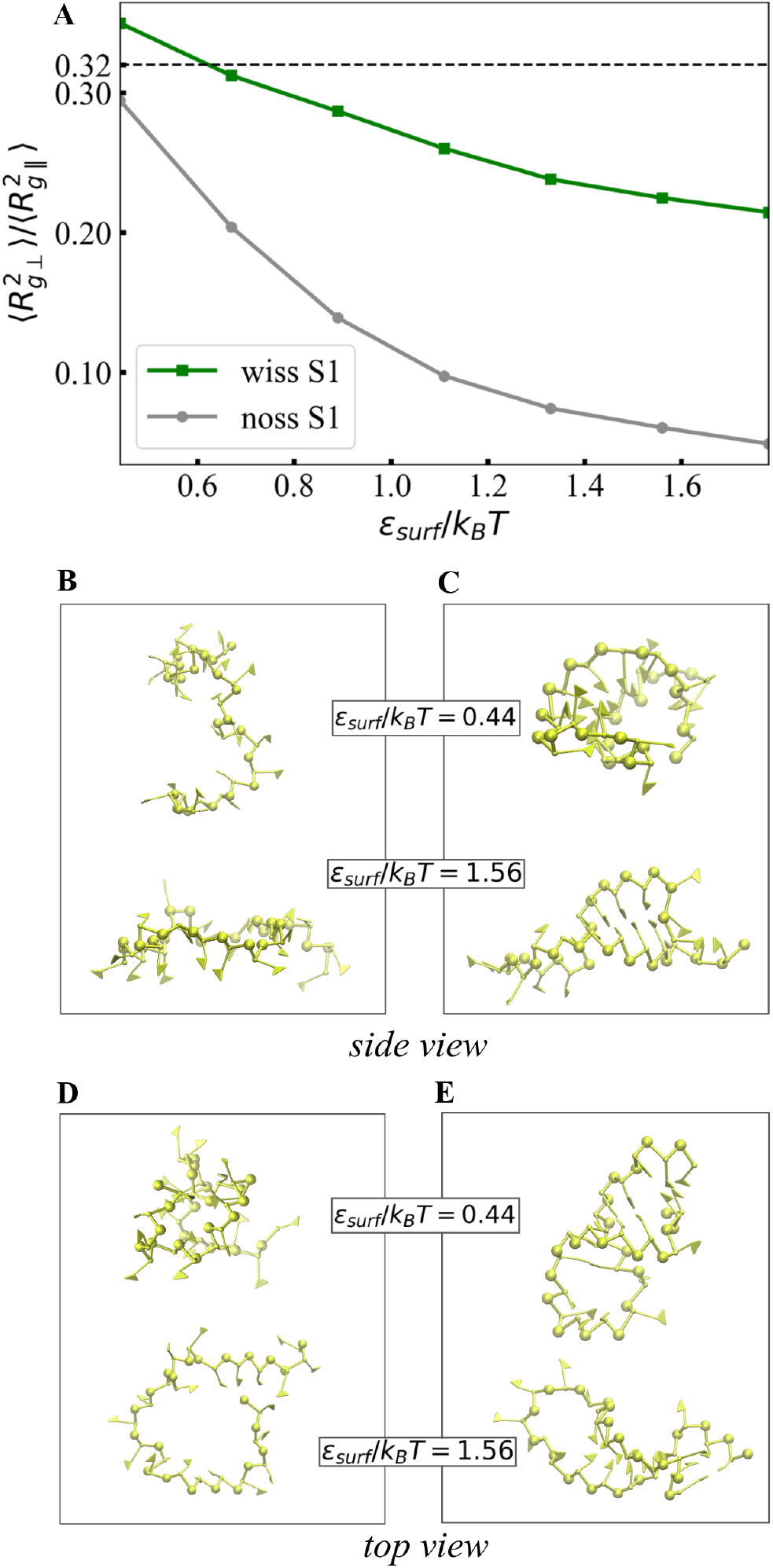
(A) Ratio of the normal and parallel radii of gyration 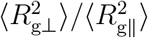 of RNA fragment S1 as a function of *ε*_surf_/*k*_B_*T*. Snapshots of the unstructured [(B) and (D)] and structured [(C) and (E)] fragments projected as side and top views with respect to the flat surface for given *ε*_surf_/*k*_B_*T* values.

### Long hairpin (RNA fragment S2)

We have performed the same analysis as above also for the longer RNA fragment, a hairpin with several additional bulges (S2-wiss), illustrated in Figure 1. This fragment includes more unpaired nucleotides than the first fragment, consequently providing a hinge-like structure of considerable flexibility, which is a factor that contributes to the structural arrangement and substrate adsorption efficiency. Interestingly, despite the differences, we observe that the two distinct regimes of adsorption behavior described before essentially persist also for the longer RNA fragment S2 (as can be seen in Figures 4 and 6).

**Figure 6:**
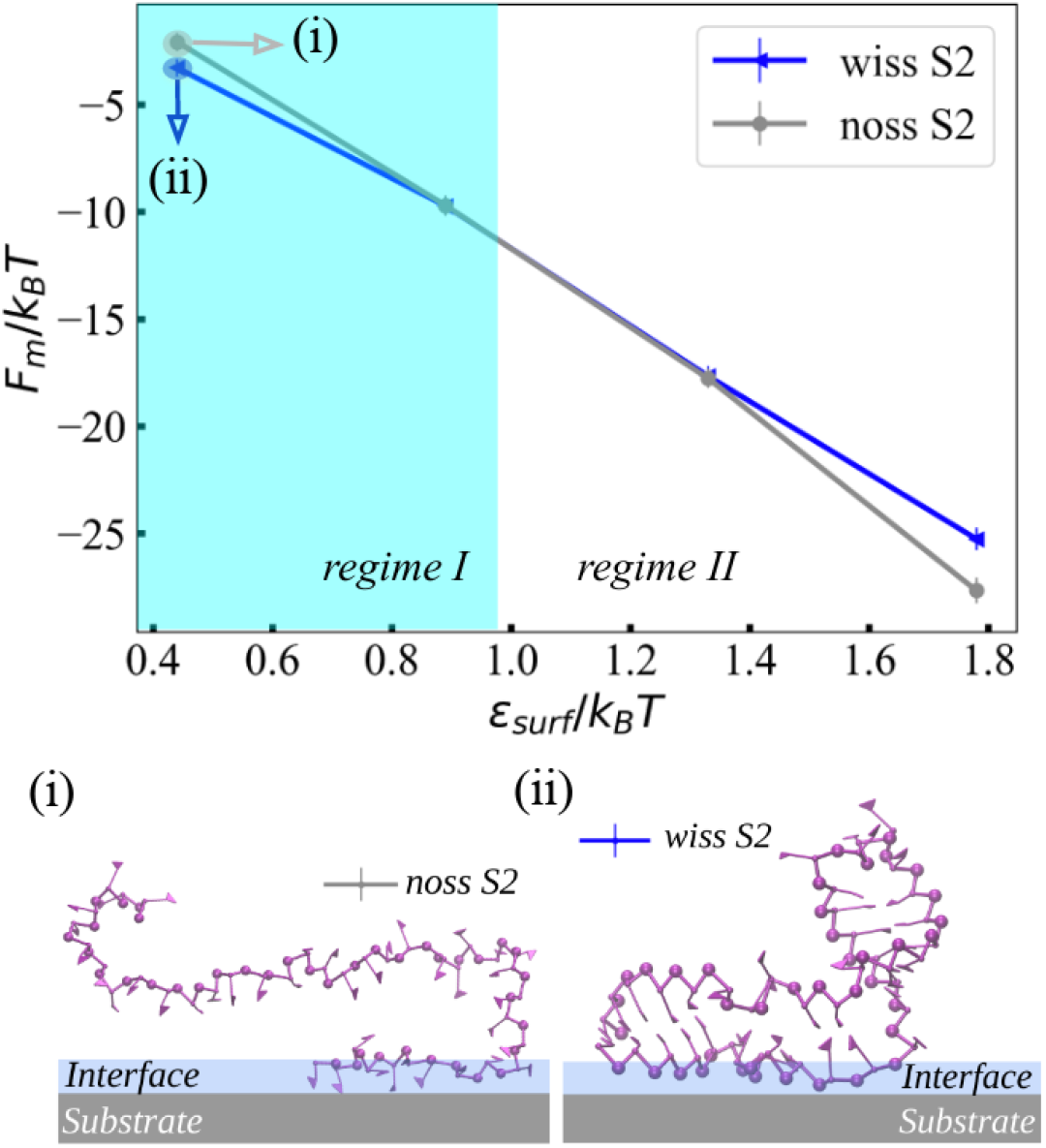
Adsorption free energy *F_m_* as a function of the substrate attraction strength *ε*_surf_ for RNA fragments S2-wiss and S2-noss. Snapshots of (i) unstructured and (ii) structured fragments at *ε*_surf_ = 0.44 *k*_B_*T*.

For the longer fragment S2, the value of *ε*_surf_ for which the interaction of the unstructured fragment S2-noss becomes more favorable than the structured S2-wiss is shifted towards larger values (*ε*_surf_/*k*_B_*T* ≈ 0.97) when compared to the smaller fragment S1. Such a shift is consistent with a point of balance between more rigid (base-paired) and more flexible (loops) regions, where, for longer unstructured molecules, a higher conformational barrier is present. The value of the transition point between the two regimes depends on the shape of the secondary structure of RNA, as seen for fragments S1 and S2. Moreover, the differences in adsorption free energies between structured and unstructured RNAs in regime I grow with the fragment size (see snapshots of the monomers trapped at the adsorption interface in Figure 6(i) and (ii)). In other words, for longer and more rigid fragments, a stronger total adsorption is expected in regime I. In regime II, however, a stronger adsorption occurs for the unstructured fragments. The resulting PMF curves are displayed in Figure S6 in Supplementary Information, while the components of the radii of gyration are listed in Tables S3 and S4.

### Multibranch fragment S3 and qualitative scaling

The last fragment with secondary structure addressed in this work is a multibranch fragment (S3-wiss), which is comprised of two hairpins joined together by a single unpaired nucleotide hinge (see Figure 1). The normalized adsorption free energy of fragment S3 as a function of *ε*_min_ is shown in Figure 7(A). Full PMF as a function of distance from the surface is shown in Figure S6, while the components of the radii of gyration are shown in Table S5. Specifically, we show that the adsorption behavior of the multibranch fragment S3, once the results are normalized for the different number of nucleotides, almost overlap with fragments S1 and S2. Here, we note that base-pair fractions are comparable for all three fragments, namely 63% (S1), 60% (S2), and 61% (S3), and we can thus conclude such an overlap only for this particular base-pair fraction.

**Figure 7:**
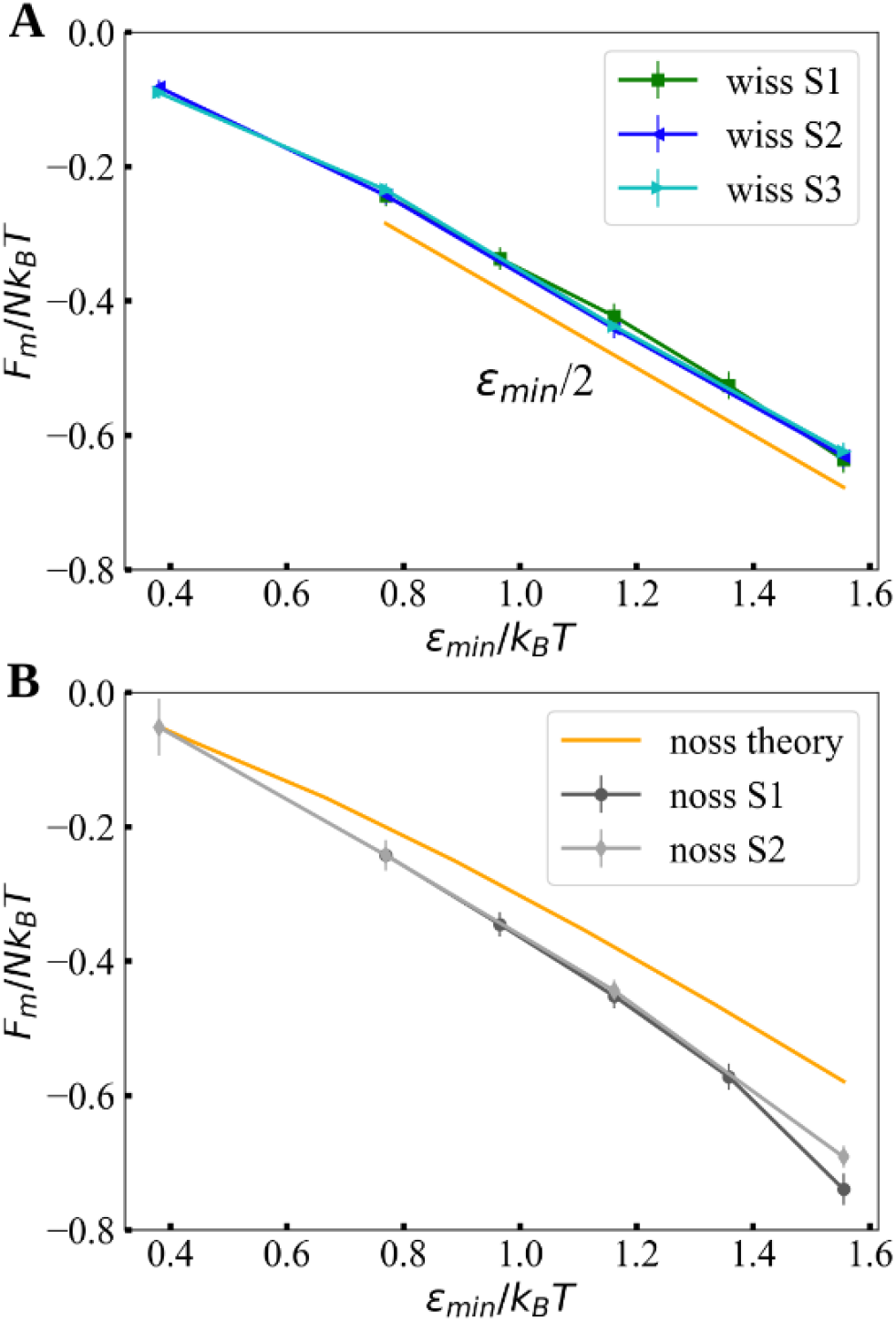
Adsorption free energy per nucleotide as a function of *ε*_min_ for the three fragments S1, S2, and S3 (A) with a secondary structure and (B) without a secondary structure. The fragments with a secondary structure show a linear tendency, shown by the slope *ε*_min_/2 (in orange). The gray curves correspond to the unstructured fragments and indicate similar trends. The orange curve in panel (B) corresponds to a one-dimensional Edwards model for a long ideal chain adsorbed onto a surface (for details, see Supplementary Information and Figure S7).

Figure 7(B) shows the comparison between two simulated unstructured RNA fragments with different lengths and the theoretical prediction based on the continuum Edwards model of an infinite ideal chain with the same adsorption potential. As seen previously in Figure 4(A) and Figure 6, the adsorption becomes more favorable as the structures grow in size. Although the difference between structured and unstructured fragments is of the order of *k*_B_*T* for low values of *ε*_min_, the adsorption properties, shown here for smaller and larger fragments, indicate that such a difference grows with the fragment size. This difference is therefore expected to become large for very long RNA fragments, for instance, in the case of the whole genome of a virus.

Moreover, for large *ε*_min_, the behavior of fragments with secondary structure resembles a straight line with a slope approximately given by *ε*_min_/2 (see Figure 7(A)). This can be rationalized by the fact that only half of the phosphates (monomers) are in direct contact with the surface because of the shape of the base-paired regions. On the contrary, fragments without secondary structure vary at a faster rate without reaching a linear regime, suggesting that more contacts may form, but the change in the shape of the molecule still plays a role in its adsorption. The result of the Edwards model for the infinitely-long ideal chain in Figure 7(B) displays essentially the same features as the ones observed in the simulations, although with a considerably larger entropy contribution, resulting in a slower variation of the free energy per monomer with the adsorption strength. Nevertheless, the decay is nonlinear, suggesting that a crossover between regimes I and II might persist, regardless of the system size.

Our results, based on the comparison between bulged-hairpin and multibranch RNA fragments show that the total fraction of base-pairs in the secondary structure determines the adsorption behavior in regime I rather than the exact topology of the RNA motifs and the additional freedom given by relative positions and orientations of the stems, at least when it comes to adsorption onto flat substrates. As for the difference of adsorption energies between structured and unstructured RNA fragments with the same sequence under regime I, it increases with the number of contacts. For instance, for a weak surface interaction with adsorption strength *ε*_surf_/*k*_B_*T*=0.44, the difference between the noss and wiss structure for the small hairpin (fragment S1) is Δ*F*_m_ ≈ 1.0 *k*_B_*T*, and for the longer bulged-hairpin (fragment S2) Δ*F*_m_ ≈ 1.5 *k*_B_*T*.

For stronger attractions in regime II, Δ*F*_m_ increases monotonically with the number of contacts (after the conformation barrier is overcome) and depends instead on the number of contacts formed with the substrate (see Figures 4 and 6 as well as Figure S5 in Supplementary Information).

## Conclusions

We have studied the role of RNA secondary structure in its adsorption to planar substrates. We use a simple and general coarse-grained RNA model interacting with the surface via a Debye-Hückel potential, which captures the basic electrostatic part of the dominant interaction in the adsorption processes on charged surfaces. Our investigation highlights the existence of two adsorption regimes concerning the RNA structure and the substrate attraction. In the first regime, operative at a weak surface attraction, base-paired segments of structured RNA behave as rigid objects and attach more easily to the substrate than unstructured, undulating RNA fragments (i.e., lacking any secondary structure). Increasing surface attraction leads to a second regime, which favors the adsorption of unstructured RNA fragments over structured ones. The existence of this turning point in adsorption does not depend on the exact nature of the interaction potential but appears to be more general. Namely, we have demonstrated that the Mie potential, based on van der Waals interaction, yields a similar behavior in adsorption. We have rationalized the origin of the two adsorption regimes as an interplay between the conformational entropy of the RNA fragments and the surface attraction. We complemented our simulations with an analytical scaling theory of the ideal chain, which provides deeper insight into the qualitative trends of adsorption of long polyelectrolytes as a function of substrate attraction strength. Based on the theoretical and simulation trends, we expect that the two observed adsorption regimes should persist even for very long RNA molecules (i.e., ≳ 1000 nucleotides in length).

Our results, which indicate a selectivity in adsorption between single- and double-stranded regions of RNA, underline the importance of RNA structure in regulating its adsorption to various substrates. We expect that the selective adsorption of one RNA structure over the other could be experimentally controlled by tuning the interaction strength, for instance, by changing the salt concentration or pH. Moreover, the lower interaction free energies of unstructured RNAs compared to structured, double-stranded ones at high attractions suggest that possibly highly attractive surfaces may promote the unfolding of a double-stranded RNA structure. This would add an additional layer of complexity to the RNA-substrate interactions, differentiating the behavior of RNA and DNA information carriers when interacting with biological substrates. These observations can have crucial and far-reaching biological implications since RNA adsorption is linked to numerous biological functions, including phosphate positioning in virus shells, stability of RNA nanocarriers, and transfection of short RNAs.^25,44,49–51^ A particularly noticeable example is RNA packaging in viral capsids, where it has been observed that unstructured RNA molecules can compete with the native RNA genome when simultaneously present in the solvent.^19,52^

In the future, it would be insightful to extend the studies to other (longer) RNA structures and different specific molecular substrate interactions and geometries. Further optimization of our code will allow us to extend and test those predictions. As for short RNA molecules, we already demonstrated that our computational model is versatile enough and generalizable to tackle more complex interaction potentials between RNA bio-macromolecules and molecular substrates.

We want to emphasize this point because several experimental techniques for secondary RNA structure characterization can be much improved by connecting them with reliable computational modeling when combined with high-throughput experimental data.^53^ RNA structure modeling combined with computational methodology is thus quickly developing and reaching higher precision/resolution, with our computational efforts fitting nicely into this newly emerging paradigm, providing additional insights into the interpretation of phenomenology as well as all the way to improving the assessment of secondary structure candidates.

## Methods

### RNA 3D structure modeling

To simulate the RNA fragments, we adopt a coarse-grained representation^54^ that offers a good balance between computational efficiency and structural detail, as shown for the 3D domain reconstruction of the STMV genome.^55^ In our model, we have implemented the potential of the flat substrate (Equations 1 and S1) and further analysis for the conformational sampling (see Supplementary Information for details). The resulting multiscale method is available in this work and hence could be further implemented in models/codes^56–60^ of preference. Figure 2 illustrates the coarse-grained representation, where each RNA nucleotide is composed of an oriented particle with a virtual site which represents its nucleoside (sugar ring plus nitrogenous base) and of a point particle which represents the phosphate group. The interactions present in the simulations are built based on an X-ray structure database, which are the backbone connectivity (through bonds and angles) and excluded volume, while the structure of the stems is enforced by an energy restraint. The restraints take on a form of harmonic potential and are applied to the nucleotides belonging to a stem in such a way as to minimize the 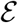RMSD^14^ metric with respect to a structure template, which is referenced to the A-form. The definition of the metric and the parameters used can be found in Supplementary Information.

### RNA simulations of adsorption and sampling analysis

The potential of mean force (PMF) was obtained systematically by means of Umbrella Sampling^61^ Monte Carlo simulations together with the Weighted Histogram Analysis Method (WHAM).^62^ The employed Collective Variable (CV) was the minimum distance between the center of mass of the RNA molecule and the surface. In this specialized procedure, each distance is consistently sampled by an individual simulation using a harmonic restraint, which improves sampling and convergence of the PMF. Histograms of the distance *d*_CM_ were used to define the PMF,

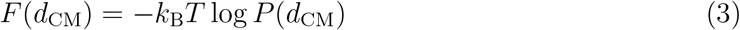

using a well-known WHAM implementation.^63^ The error bars were estimated by the bootstrapping method after carefully determining the auto-correlation time of each Monte Carlo simulation by means of blocking analysis. Additional parameters of the simulations can be found in Supplementary Information. Tables S6, S7, and S8 in Supplementary Information contain details on the CV constraint parameters of each run. Histograms of the CV are given in Figures S8 and S9 in Supplementary Information, which highlight the quality of the systematic sampling used in our calculations. Further simulation scripts and analysis files can be found at https://zenodo.org/record/4646934.

## Abbreviations

RNA: 
PMF: 
STMV: 
wiss: 
noss: 
3D: 

## Acknowledgement

H.V.G., A.B., and M.K. acknowledge financial support from the Slovenian Research Agency ARRS (Funding No. P1-0055 and J1-1701). S.P. acknowledges the project FONDECYT Iniciación en Investigación 11181334 for financial support.

## Supporting Information Available

Detailed statistical and computational details can be found in Supplementary Information. Animated simulation snapshots for each of the RNA fragments are also provided in the Supplementary Information.

