## Supplemental material for "RNA secondary structure regulates fragments’ adsorption onto flat substrates"

#### 1 Adsorption free energies for the Mie potential

In addition to the Debye-Hückel potential, we also performed calculations with the Mie 9-3 potential for the interaction between the surface and the phosphates. The Mie interaction is given by

$$U_{\text{surf}}(z) = \epsilon_{\text{surf}} \left[ \left(\frac{\sigma}{z}\right)^9 - \left(\frac{\sigma}{z}\right)^3 \right] \tag{S1}$$

where $z$ is the distance between the phosphate bead and the substrate. The interaction range is on the order of $\sigma = 0.1$ nm, which is substantially shorter than in the Debye-Hückel case, where it is $\lambda_{\rm D} = 1$ nm. The strength of the attraction was adjusted by the parameter $\epsilon_{\rm surf}$, which ranged from 6.22 to 8.89 $k_{\rm B}T$ in order to observe the adsorbed regime for both noss and wiss scenarios. Additional details concerning the length and Umbrella Sampling implementation of the simulations are in the Simulation Parameters section of the Supplementary Information.

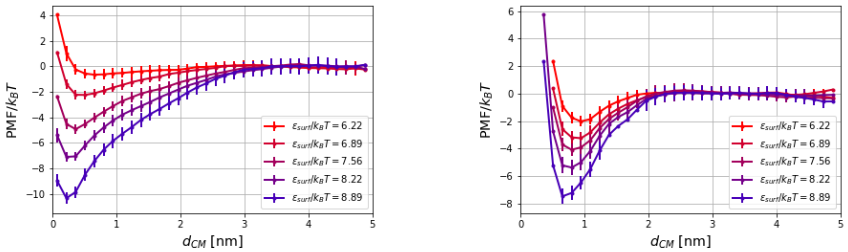


Figure S1: PMF curves for RNA fragments S1-noss (left panel) and S1-wiss (right panel) for RNA-substrate interaction using the Mie potential.

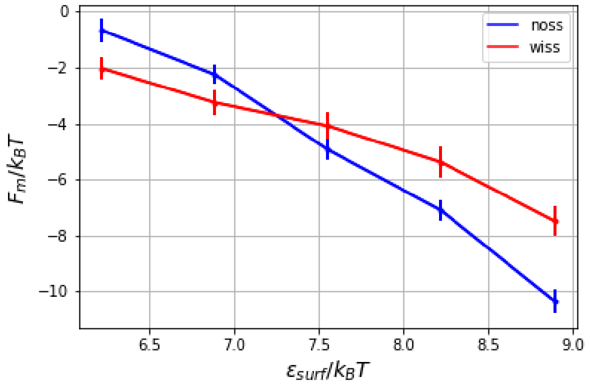


Figure S2: Minimum of PMF for RNA fragment S1 using the Mie potential for RNA-substrate interaction.

PMF curves for RNA fragment S1 in both noss and wiss forms are shown in Figure S1, and the minimum values of the PMF are depicted in Figure S2, displaying the same trend as in the case of the Debye-Hückel potential, analyzed in the main text. In Figure S2, we identify the same two adsorption regimes as in the main text, the first for $\epsilon_{\rm surf} < 7.25$, where the fragment with a secondary structure adsorbs more strongly, and the second regime for stronger adsorption strengths ($\epsilon_{\rm surf}$ >7.25), in which the unstructured molecule shows a more favorable adsorption. These results are comparable to those shown in Figure 4 and discussed in the main part of the article.

#### 2 Monomer distributions

Figure S3 shows the monomer distribution for RNA fragments S1-wiss and S1-noss, considering only those conformations where the molecule's center of mass is at a distance equal to or less than $d_m$, where $d_m$ is the point at which PMF has half of the value of its minimum. For RNA fragments S2 and S3, histograms showing their monomer distributions are shown in Figure S4.

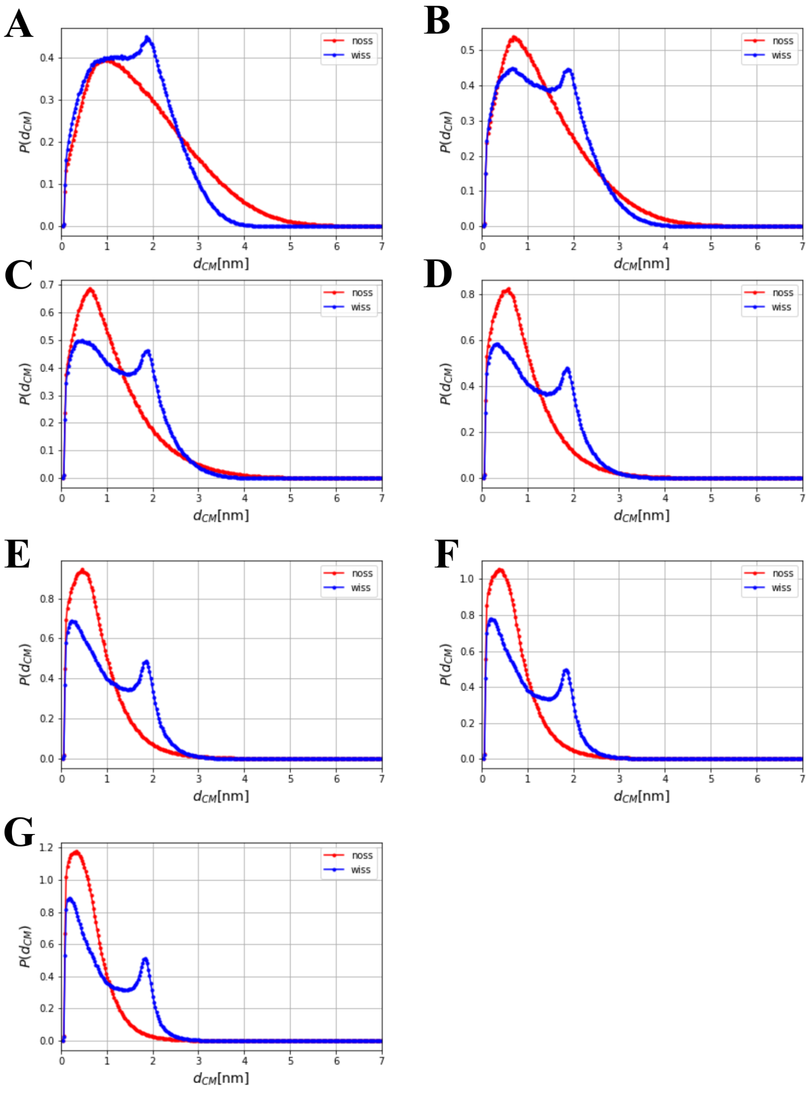


Figure S3: Monomer (phosphate group) distribution for adsorbed structures as a function of the surface distance of RNA fragments S1-noss (red lines) and S1-wiss (blue lines), for values of $\epsilon_{\text{surf}}/k_BT$ of (a) 0.44, (b) 0.67, (c) 0.89, (d) 1.11, (e) 1.33, (f) 1.56, and (g) 1.78.

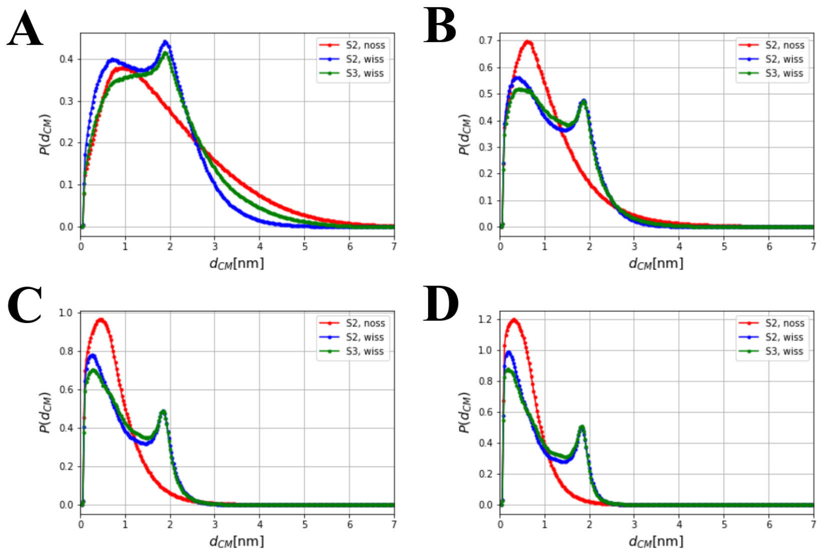


Figure S4: Monomer (phosphate group) distribution for adsorbed structures as a function of the surface distance of RNA fragments S2 (in noss and wiss forms) and S3-wiss, for values of $\epsilon_{\mathrm{surf}}/k_B T$ of (a) 0.44, (b) 0.89, (c) 1.33, and (d) 1.78.

#### 3 Number of contacts for RNA fragments S2 and S3

In Figure S6, we show the number of contacts for RNA fragments S2 and S3. The blue (S2-noss) and red (S2-wiss) curves indicate a similar behavior of the number of contacts with adsorption strength as shown in Figure 4B in the main text for RNA fragment S1. Additionally, the number of contacts for fragment S3-wiss is expected to be equal to or greater than for S2-wiss.

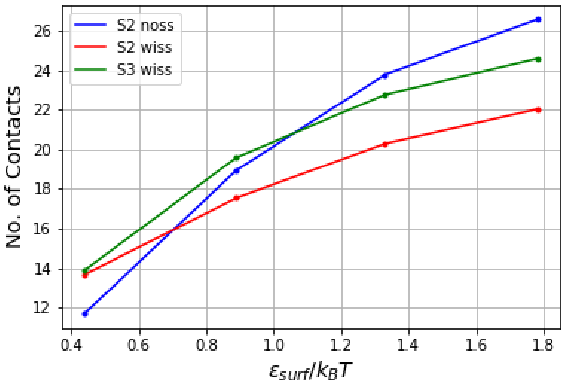


Figure S5: Number of contacts of RNA fragments S2 and S3 on the adsorbing surface interacting via the Debye-Hückel potential.

#### 4 Radius of Gyration components

Considering that the number of coarse-grained particles is $N$ and that the attractive surface is perpendicular to the $z$ direction, the definitions of the normal $\langle R^2_{\mathrm{g}\perp} \rangle$ and parallel $\langle R^2_{\mathrm{g}\parallel} \rangle$ contributions to the radius of gyration are given by

$$\langle R^2_{\mathrm{g}\perp} \rangle = \left\langle \frac{1}{N} \sum_i \left[ (x_i - x_{\mathrm{CM}})^2 + (y_i - y_{\mathrm{CM}})^2 \right] \right\rangle \tag{S2}$$

$$\langle R^2_{\mathrm{g}\parallel} \rangle = \left\langle \frac{1}{N} \sum_i (z_i - z_{\mathrm{CM}})^2 \right\rangle , \tag{S3}$$

where $\mathbf{R}_{\mathrm{CM}} = (x_{\mathrm{CM}}, y_{\mathrm{CM}}, z_{\mathrm{CM}})$ is the position of the center of mass. Tables S1 and S2 list the components of the radius of gyration for RNA fragment S1 in both its structured and unstructured variant, while Tables S3, S4, and S5 contain the components of the radius of gyration for S2-noss, S2-wiss, and S3-wiss RNA fragments, respectively.

| $\epsilon_{\text{surf}}$ [$k_B$T] | $\langle R^2_{g,\parallel} \rangle$ [$nm^2$] | $\langle R^2_{g,\perp} \rangle$ [$nm^2$] |
|---|---|---|
| 0.44 | 3.23 | 0.95 |
| 0.67 | 3.58 | 0.73 |
| 0.89 | 3.79 | 0.52 |
| 1.11 | 4.05 | 0.39 |
| 1.33 | 4.30 | 0.32 |
| 1.56 | 4.43 | 0.26 |
| 1.78 | 4.59 | 0.22 |

Table S1: Radius of gyration components for S1-noss.

| $\epsilon_{\text{surf}}$ [$k_B$T] | $\langle R^2_{g,\parallel} \rangle$ [$nm^2$] | $\langle R^2_{g,\perp} \rangle$ [$nm^2$] |
|---|---|---|
| 0.44 | 1.46 | 0.51 |
| 0.67 | 1.51 | 0.47 |
| 0.89 | 1.53 | 0.44 |
| 1.11 | 1.57 | 0.41 |
| 1.33 | 1.6 | 0.38 |
| 1.56 | 1.62 | 0.36 |
| 1.78 | 1.65 | 0.35 |

Table S2: Radius of gyration components for S1-wiss.

| $\epsilon_{\text{surf}}$ [$k_B$T] | $\langle R^2_{g,\parallel} \rangle$ [$nm^2$] | $\langle R^2_{g,\perp} \rangle$ [$nm^2$] |
|---|---|---|
| 0.44 | 7.67 | 1.55 |
| 0.89 | 9.12 | 0.6 |
| 1.33 | 10.73 | 0.3 |
| 1.78 | 11.47 | 0.21 |

Table S3: Radius of gyration components for S2-noss.

| $\epsilon_{\text{surf}}$ [$k_B$T] | $\langle R^2_{g,\parallel} \rangle$ [$nm^2$] | $\langle R^2_{g,\perp} \rangle$ [$nm^2$] |
|---|---|---|
| 0.44 | 3.69 | 0.58 |
| 0.89 | 4 | 0.42 |
| 1.33 | 4.19 | 0.34 |
| 1.78 | 4.35 | 0.31 |

Table S4: Radius of gyration components for S2-wiss.

### 5 Potential of Mean Force of RNA fragments S2 and S3

The Collective Variable (CV) restraint consisted on a harmonic potential whose minimum was situated at a distance $d_0$, which ranged between 0.5 nm and

| $\epsilon_{\text{surf}}$ [k$_B$T] | $\langle R^2_{g,\parallel} \rangle$ [nm$^2$] | $\langle R^2_{g,\perp} \rangle$ [nm$^2$] |
|---|---|---|
| 0.44 | 4.34 | 0.79 |
| 0.89 | 4.82 | 0.45 |
| 1.33 | 5.02 | 0.35 |
| 1.78 | 5.08 | 0.32 |

Table S5: Radius of gyration components for S3-wiss.

5.5 nm for S1-wiss and between 0.5 nm and 6.5 nm for S1-noss, with a spacing of 0.5 nm. The values of $\varepsilon_{\text{surf}}/k_{\text{B}}T$ were first determined in SPQR reduced units and further converted (as shown in the Simulation Parameters section). For systems S2 and S3, the CV restraints were located between 0.5 nm and 8 nm, with a spacing of 0.5 nm, and the surface strength was determined for four values as in the case of fragment S1. For each Umbrella Sampling simulation restraint, 24 independent simulations were performed.

The PMFs were shifted to zero at the positions where their centered derivatives were consistent with zero within the error bar, which is between 4.5 and 5 nm for S1-wiss and between 5.5 and 6 nm for S1-noss. For fragments S2 and S3, this domain was set between 7 and 8 nm. For the rest of the averages, the same weights were used on a sampled subspace of conformations. The average radius of gyration was calculated on a subset of structures with a phosphate group at a distance from the surface below 0.37 nm, approximately one phosphate radius away from the surface potential minimum, as defined within the model. In this analysis, the calculation of the number of contacts considered only snapshots whose value of $d$ was below $d_{\text{m}}$, the distance at which the PMF reaches half of its minimum value. Figure S6 show the PMF of the RNA fragments S2 and S3 .

### 6 Free energy per monomer for an ideal adsorbed chain

We briefly describe the basics of the formalism that we have used to estimate the free energy per monomer of an adsorbed ideal polymer chain composed of $N$ monomers and with a bond length $l$. In our case, bond length is taken as the average distance between two phosphate groups of an independent simulation of RNA fragment S1-noss in the absence of a surface, resulting in $l = 0.61$ nm. If the probability of finding the first monomer at position $\mathbf{r}_0$ is $P(\mathbf{r}_0; 0)$, the probability of finding its $N$-th unit at a position $\mathbf{r}$ can be expressed as

$$P(\mathbf{r}; N) = \int d\mathbf{r}_0 G_0(\mathbf{r}, \mathbf{r}_0; N) P(\mathbf{r}_0; 0). \tag{S4}$$

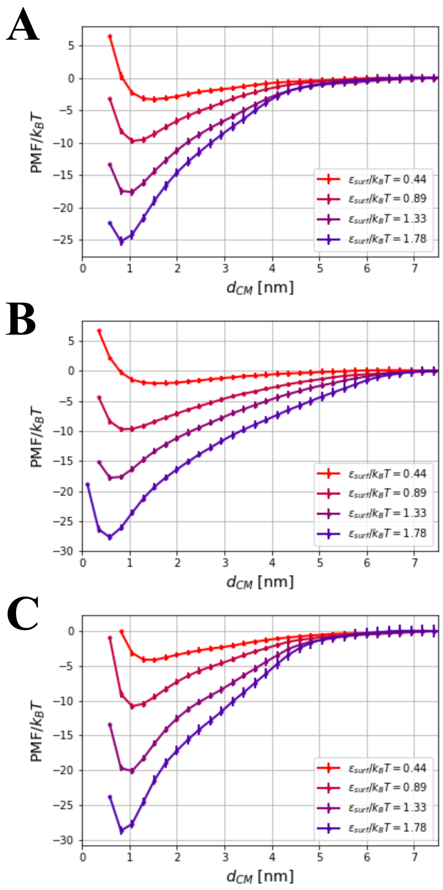


Figure S6: PMF of fragments (A) S2-noss, (B) S2-wiss, and (C) S3-wiss.

Here, $G_0(\mathbf{r}, \mathbf{r}_0; N)$ is the polymer's Green function, which for an ideal chain satisfies the Edwards equation [4]

$$-\frac{\partial}{\partial N} G(\mathbf{r}, \mathbf{r}_0; N) = \left[ -\frac{l^2}{6} \nabla^2 + \beta u(\mathbf{r}) \right] G(\mathbf{r}, \mathbf{r}_0; N), \tag{S5}$$

where $u(\mathbf{r})$ is an external potential exerted on the monomers.

The Edwards equation can be solved by writing an Ansatz of the form [4]

$$G(\mathbf{r}, \mathbf{r}_0; N) = \sum_{n=0}^{\infty} e^{-N\epsilon_n} \psi_n(\mathbf{r}) \psi_n(\mathbf{r}'), \tag{S6}$$

considering that $G(\mathbf{r}, \mathbf{r}_0; 0) = \delta(\mathbf{r} - \mathbf{r}_0)$. The values of $\epsilon_n$ introduced on the right-hand side are the eigenvalues of the linear differential operator $\mathcal{L}_E = -\frac{l^2}{6}\nabla^2 + \beta u(\mathbf{r})$. In addition, the functions $\psi_n(\mathbf{r})$ form a complete and orthonormal basis for the space of solutions given by the properties of $\mathcal{L}_E$. For large $N$, the expression of $G$ can be approximated by

$$G(\mathbf{r}, \mathbf{r}_0; N) \approx e^{-N\epsilon_0} \psi_n(\mathbf{r}) \psi_n(\mathbf{r}') \tag{S7}$$

where $\epsilon_0$ is the smallest eigenvalue.

Several important quantities can be obtained from the solution of the previous equation in terms of its eigenfunctions and eigenvalues. The most relevant for our purposes is the partition function $Z_N$, written as

$$Z_N \propto \int d\mathbf{r}_0 \int d\mathbf{r}\, G(\mathbf{r}_0, \mathbf{r}; N). \tag{S8}$$

up to a multiplicative constant, from which the free energy can be obtained under the present assumptions as

$$F(N) = -k_{\mathrm{B}} T \log Z_N \approx N k_{\mathrm{B}} T \epsilon_0 \tag{S9}$$

plus an additive constant independent of $N$. In addition, the monomer concentration at $\mathbf{r}$ is given by

$$c(\mathbf{r}) \approx N \psi_0(\mathbf{r})^2. \tag{S10}$$

Considering an infinite surface, the potential depends only on the direction $x$, yielding an external potential $u(x)$, and we have to solve the equation in one dimension. The eigenvalue $\epsilon_0$, according to Equation S9 , is the free energy per monomer divided by $k_{\mathrm{B}}T$, which has a negative value in the adsorbed regime. The Edwards equation adopts the form

$$\left[ -\frac{l^2}{6} \frac{\partial^2}{\partial x^2} + \beta u(x) \right] \psi_0(x) = \epsilon_0 \psi_0(x) \tag{S11}$$

with the boundary condition $\psi_0(0) = \psi_0(x \to \infty) = 0$. The free energy per monomer is

$$\frac{F}{N} = \int_0^{\infty} dx\, u(x) \psi_0(x)^2 - \frac{l^2 k_{\mathrm{B}} T}{6} \int_0^{\infty} dx\, \psi_0(x) \frac{d^2}{dx^2} \psi_0(x), \tag{S12}$$

where the first and second terms of the right-hand side can be interpreted as the surface energy and the entropy loss due to the adsorption process, respectively. It is worth pointing out that the surface energy term resembles the number of contacts as defined in the main text.

To compare this result with our simulations, we assume $u(x) = -u_w \exp(-x/\lambda)$. The repulsive part is taken into account via a boundary condition and is not explicitly included in the above expression. Therefore, for comparison, we interpret $u_w$ as the minimum value of the surface potential used in the simulations. The most general solution of the one-dimensional Edwards equation which is consistent with the boundary conditions is

$$\psi_0(x) = C J_\alpha \left( \frac{2\sigma}{l} \sqrt{6\beta u_w} e^{-x/2\lambda} \right), \tag{S13}$$

where $J_\alpha$ is the Bessel function of the first kind and order $\alpha$, $C$ is a constant which can be obtained by normalizing the monomer concentration, and $\alpha = 2\sigma\sqrt{6|\epsilon_0|}/l$. Additionally, the boundary conditions require that the order of the Bessel function has to be greater than zero, while the eigenvalue $\epsilon_0$ must be determined numerically.

The free energy per monomer, together with its energy and entropy contributions, are shown in Figure S7 for a range of $u_w$ and $\beta = 1$. The entropy contribution $-TS/N$ is positive and increases with $u_w$, since stronger attraction increases the polymer confinement and reduces the absolute value of $S$. On the other hand, the energy term, proportional to the number of contacts, decays monotonically—almost linearly in the shown regime. Interestingly, in this model, the entropy has a larger contribution to the free energy than in the simulated systems. Clearly, this is an overestimation compared to our simulations, where the excluded volume interactions restrict the conformation space of the RNA chain.

### 7 Definition of secondary structure restraints

Structural restraints minimize the $\mathcal{E}$RMSD metric [1] between a stem and a template stem constructed with standard parameters of an A-form RNA. This metric is defined in terms of the set of vectors $\{\,\mathbf{r}_{ij}\}$ and $\{\,\mathbf{r}^{\mathrm{r}}_{ij}\}$ obtained from the simulated and template structures, respectively, calculated from the position of nucleobase $i$ with respect to a reference frame situated at the origin of nucleobase $j$ with its orientation. The set of rescaled vectors $\tilde{\mathbf{r}} = (r_x/a, r_y/b, r_z/c)$, with $a = b = 0.5$ nm and $c = 0.3$ nm, is used to define the vectors $\mathbf{G}(\tilde{\mathbf{r}}_{ij}) = (\sin(\gamma\tilde{r}_x/\tilde{r}), \sin(\gamma\tilde{r}_y/\tilde{r}), \sin(\gamma\tilde{r}_z/\tilde{r}), 1 + \cos(\gamma\tilde{r}))\,\Theta(\tilde{r}_{\mathrm{c}} - \tilde{r})/\gamma$, where $\gamma = \pi/\tilde{r}_{\mathrm{c}}$, $\Theta$ is the Heaviside function and $\tilde{r}_{\mathrm{c}}$ is a cutoff parameter. For a particular fragment (or stem) denoted by $s$ and composed of $N_s$ nucleotides, the $\mathcal{E}$RMSD can be written

$$E_s = \sqrt{\frac{1}{N_s} \sum_{j,k} \left| \mathbf{G}(\tilde{\mathbf{r}}_{jk,\mathrm{s}}) - \mathbf{G}(\tilde{\mathbf{r}}^{\mathrm{r}}_{jk,\mathrm{s}}) \right|^2} \tag{S14}$$

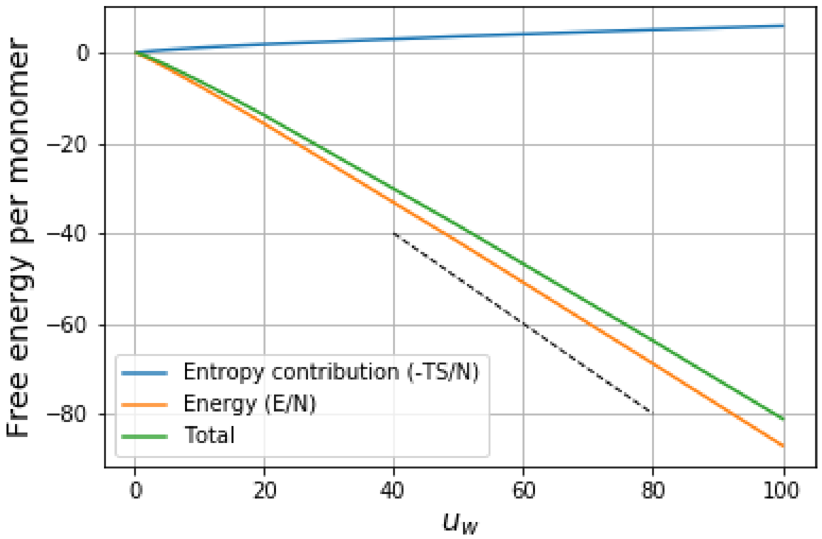


Figure S7: Free energy per monomer for an ideal chain, separately showing also the energy $E/N$ and entropy $-TS/N$ contributions.

which allows us to define the restraining potential as

$$U_{\mathrm{ss}} = \frac{K_{\mathrm{ss}}}{2} \sum_s E_s^2 \tag{S15}$$

where $K_{\mathrm{ss}}$ is the corresponding harmonic spring constant [1]. In this manner, the relative positions and orientations of the nucleobases belonging to each stem are enforced to have the values of a reference A-form stem. This procedure has also been used for enforcing other motifs and backmapping RNA structures predicted by SPQR into an all-atom representation by means of Steered-Molecular Dynamics [2].

### 8 Simulation Parameters

We choose the temperature to be $T = 9u_{\mathrm{cg}}$ for all the simulations listed here, where $u_{\mathrm{cg}}$ is the SPQR energy unit, in order to obtain a better resemblance of the distribution functions sampled by the backbone potentials, originally obtained from an X-ray structure set [3]. This choice is a technicality that does not affect the results in practice because all the bonded interactions are essentially unaffected by this parameter, and all non-bonded interactions (apart from the excluded volume) are incorporated through the secondary structure restraint. Thus, the simulated scenarios are below the melting temperature of the structured fragments, where unstructured chains of the same length can coexist.

In addition, both the glycosidic bond angle state and the sugar pucker were kept fixed to their *anti* and $C3'endo$ conformations during the calculations to avoid the introduction of additional parameters. In these units, the chosen values of $\epsilon_{\rm surf}$ described in the main text have values of $4u_{\rm cg}$, $6u_{\rm cg}$, $8u_{\rm cg}$, $10u_{\rm cg}$, $12u_{\rm cg}$, $14u_{\rm cg}$, and $16u_{\rm cg}$. This range of values of $\epsilon_{\rm surf}$ allows us to clearly distinguish the two adsorption regimes described in the main text. Secondary structure was fixed with a harmonic restraint on the $\mathcal{E}$RMSD, with an dimensionless cutoff $\tilde{r} = 100$ and a harmonic spring of $K_{ss} = 50\epsilon_{\rm cg}$.

The RNA fragments S1, S2, and S3 with secondary structure were constructed imposing the secondary structure conditions until the contacts were formed, relaxed with different random seeds and integrated for $10^6$ Monte Carlo (MC) sweeps for the fragment S1 and $10^8$ MC sweeps for the fragments S2 and S3. Each MC sweep consists of $N$ MC steps, where $N$ is the number of nucleotides. From here, the unstructured fragments were generated for each initial condition, running a simulation at constant temperature for $10^6$ MC sweeps. It was noticed that starting from a helical conformation, as a single strand extracted from a template stem in the A-form, gave a consistent end-to-end distance after the relaxation procedure. For systems S2 and S3, the procedure was the same, although the equilibration simulations for decorrelating the structured fragments and generating the unstructured initial conditions consisted of $10^8$ MC sweeps. After this, the surface was positioned perpendicular to the $z$ direction, 0.1 nm from the closest nucleotide. The simulations were performed for $1.5 \times 10^8$ MC sweeps, and configurations were saved every 50000 steps, while the first 750 saved configurations were discarded. For fragments S2 and S3, the runs consisted of $3 \times 10^8$ MC sweeps, and configurations were saved every 10000 steps while the first 750 saved configurations were again discarded.

The strength of the harmonic spring used to fix the RNA center-of-mass to a chosen value of $d_0$ at each Umbrella Sampling simulation was $k_{\rm u} = 5.56\ k_{\rm B}T/{\rm nm}^2$. In order to improve the convergence and statistics, a value of 11.12 $k_{\rm B}T/{\rm nm}^2$ was used in some cases, which are specified in Table S6 together with any simulations added for improving the convergence or removed in case they did not contribute to the statistics. The unweighted distributions are plotted in Figures S8 and S9 for S1-noss and S1-wiss, respectively.

| Simulation ($\epsilon_{\mathrm{surf}}/k_{\mathrm{B}}T$) | Removed<br>$d_0$/nm, $k_u$nm$^2/k_BT$ | Added<br>$d_0$/nm, $k_u$nm$^2/k_BT$ |
|---|---|---|
| S1-wiss (1.56) | 0.5, 11.12 | - |
| S1-wiss (1.78) | 0.5, 11.12 | - |
| S1-wiss (1.78) | 1, 11.12 | - |
| S2-noss, (1.78) | 3, 5.56 | 2, 11.12<br>2.5, 11.12<br>3, 11.12 |
| S2-wiss, (1.78) | 3.5, 5.56 | 2, 11.12<br>2.5, 11.12<br>3, 11.12 |
| S3-wiss, (1.56) | 3, 5.56 | 2, 11.12<br>2.5, 11.12<br>3, 11.12 |
| S3-wiss, (1.56) | 3.5, 5.56 | 2, 11.12<br>2.5, 11.12<br>3, 11.12 |

Table S6: Umbrella Sampling simulation details for all fragments under the effect of the Debye-Hückel potential.

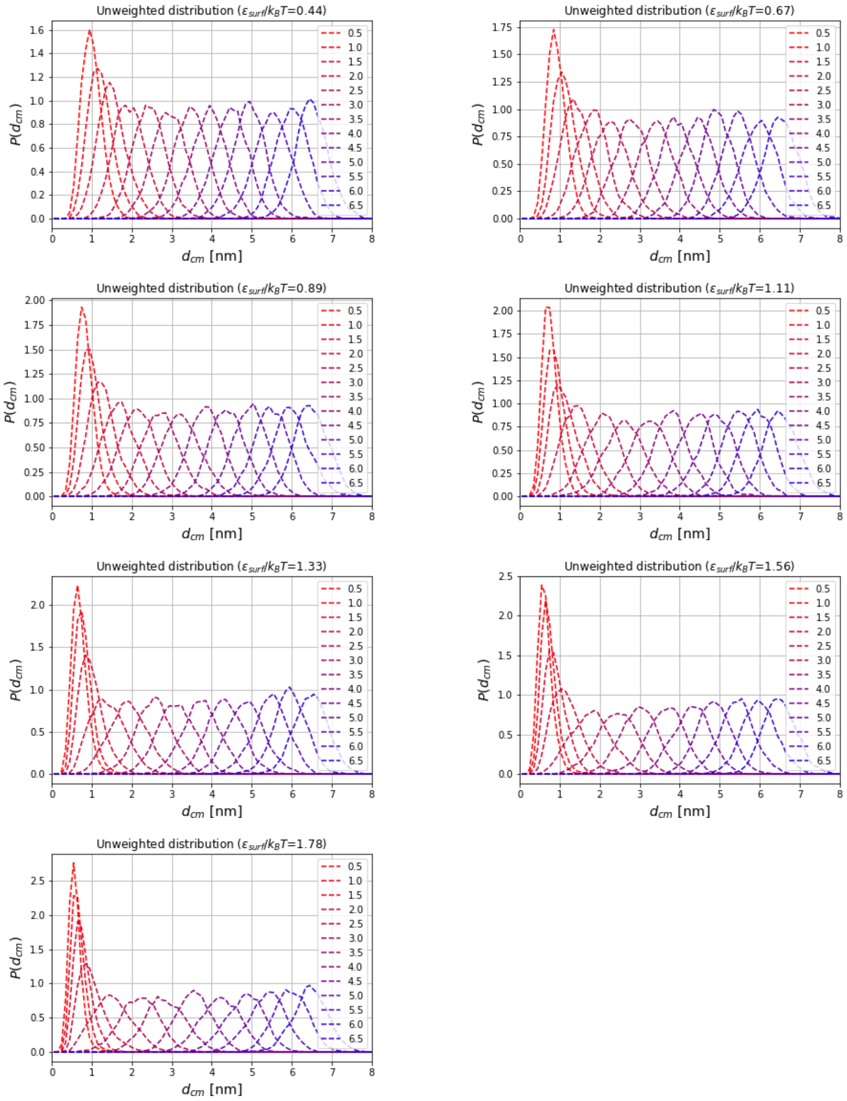


Figure S8: Unweighted distributions for fragment S1-noss.

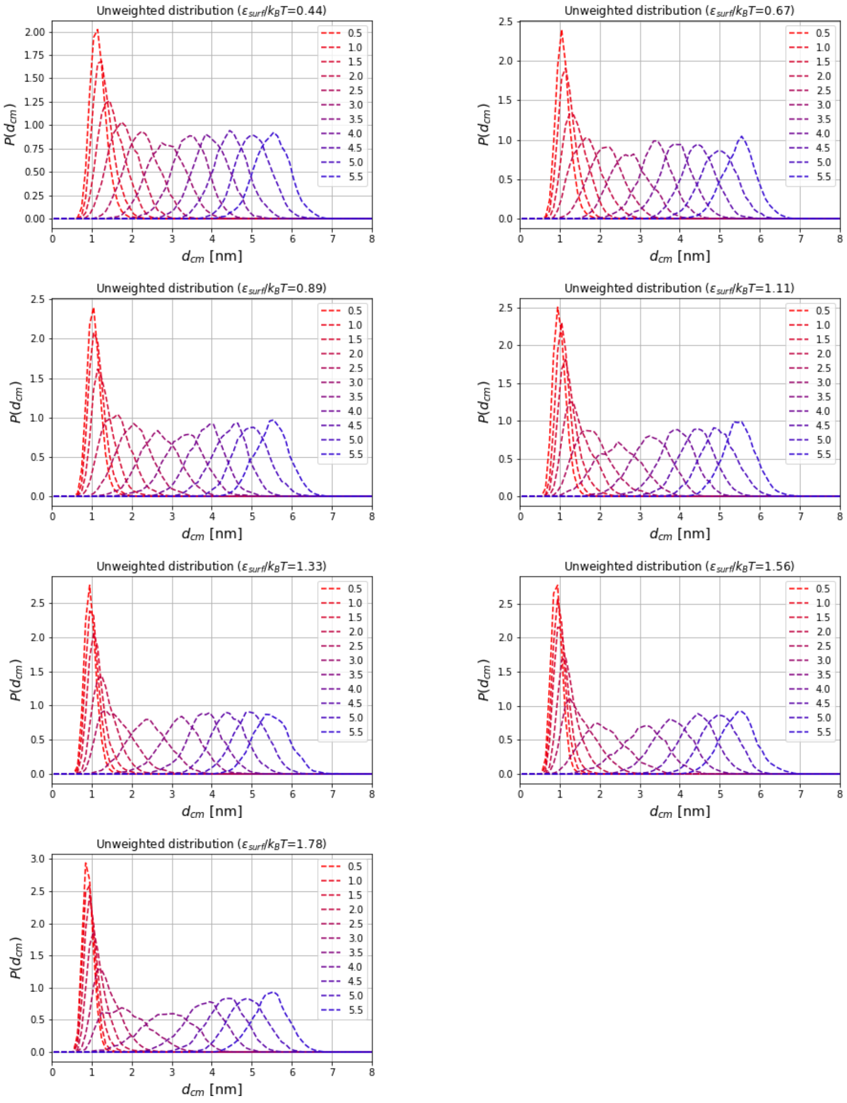


Figure S9: Unweighted distributions for fragment S1-noss.

For the simulations with the Mie potential, the harmonic restraints of the Umbrella simulations were situated at $d_0 = 0.3$ nm and from 0.5 to 4 nm, with steps of 0.5 nm and $k_u = 5.56\ \mathrm{nm}^2/k_\mathrm{B}T$. As in the previous system, in some cases, additional simulations using different Umbrella restraints were performed or omitted because of their long autocorrelation times, which are described in Table S7. For each setup, 24 independent runs were analyzed, consisting of $150 \times 10^6$ MC sweeps, and configurations were saved every 50000 sweeps. The first 750 saved configurations were discarded.

| Simulation( $\epsilon_\mathrm{surf}/k_\mathrm{B}T$) | Removed $d_0$/nm, $k_u\mathrm{nm}^2/k_\mathrm{B}T$ | Added $d_0$/nm, $k_u\mathrm{nm}^2/k_\mathrm{B}T$ |
|---|---|---|
| S1-wiss (6.22) | 2, 11.12 | 2, 5.56 |
| S1-wiss (6.89) | 2, 5.56<br>0.3, 5.56 | 2, 11.12 |
| S1-wiss (7.56) | 2, 5.56<br>2.5, 5.56 | 2, 11.12<br>2.5, 11.12 |
| S1-wiss (8.89) | 2.5, 5.56<br>0.3, 5.56 | 2.5, 11.12 |

Table S7: Details of Umbrella Sampling simulations for fragments S1-wiss and S1-noss interacting with the Mie potential.

Scripts and analysis files can be found at https://zenodo.org/record/4646934.

[4] Wokyung Sung. *Statistical Physics for Biological Matter*. Springer, 2018.
